# Predicting host-based, synthetic lethal antiviral targets from omics data

**DOI:** 10.1101/2023.08.15.553430

**Authors:** Jeannette P. Staheli, Maxwell L. Neal, Arti Navare, Fred D. Mast, John D. Aitchison

## Abstract

Traditional antiviral therapies often have limited effectiveness due to toxicity and development of drug resistance. Host-based antivirals, while an alternative, may lead to non-specific effects. Recent evidence shows that virus-infected cells can be selectively eliminated by targeting synthetic lethal (SL) partners of proteins disrupted by viral infection. Thus, we hypothesized that genes depleted in CRISPR KO screens of virus-infected cells may be enriched in SL partners of proteins altered by infection. To investigate this, we established a computational pipeline predicting SL drug targets of viral infections. First, we identified SARS-CoV-2-induced changes in gene products via a large compendium of omics data. Second, we identified SL partners for each altered gene product. Last, we screened CRISPR KO data for SL partners required for cell viability in infected cells. Despite differences in virus-induced alterations detected by various omics data, they share many predicted SL targets, with significant enrichment in CRISPR KO-depleted datasets. Comparing data from SARS-CoV-2 and influenza infections, we found possible broad-spectrum, host-based antiviral SL targets. This suggests that CRISPR KO data are replete with common antiviral targets due to their SL relationship with virus-altered states and that such targets can be revealed from analysis of omics datasets and SL predictions.

## INTRODUCTION

We recently proposed that synthetic lethality, a well-established concept in cancer therapy (1–11), could mitigate multiple known shortcomings of current antiviral drugs such as the paucity of options, restricted target spectrum due to the small genome size of viruses, the often rapid development of drug resistance (12), marginal effectiveness due to dose-limiting toxicity (13), and the necessity to redesign drugs as new viral strains emerge (14, 15). Of 92 approved antiviral drugs in 2018, two-thirds were aimed at HIV and HCV and almost 90% were small molecule drugs (16, 17). A mere 10% were host-based therapeutics, half of which were interferon-related biologics. Given the potential advantages of host-based drugs, there is a pressing need for novel host protein targeting methodologies and associated target prediction strategies (18, 19).

In the context of viral infections, synthetic lethality describes the vulnerability of virus-induced cellular states to host-based drug intervention (20). As obligate intracellular pathogens, viruses require the host cell machinery for every step of the viral life cycle including attachment, penetration, uncoating, gene expression and replication, assembly, and finally, virion release (21). During these processes, viruses effectively remodel host cells, converting them into viral factories by leveraging numerous host cell functions, leading to wide-ranging cellular changes (22). These changes are initiated by the direct interaction of viral proteins with host proteins, usurping their functions which leads to a cascade of direct and indirect effects, including altered protein complexes, changes in RNA and protein abundances, protein mis-localization, as well as changes in post-translational modifications, protein cleavage, splice patterns, and metabolomic and regulatory networks, to name a few. This reliance of viruses on host processes shifts the host cell state and introduces specific vulnerabilities to infected cells.

As an antiviral strategy, synthetic lethality capitalizes on these virus-induced vulnerabilities and the dependency of cells on the integrity of genetic networks. Notably, while many genes can be disrupted individually without undermining cell survival, specific pairs of gene disruptions are lethal in combination (9, 23–26). When a virus compromises a host protein’s normal functionality, this host protein is designated a viral-induced hypomorph (VIH) (20). Synthetic lethal partners of these VIHs can be strategically targeted to impede the viral production machinery, cellular viability, or both, providing a means of selectively disrupting the viral lifecycle to contain viral dissemination and disease, while sparing uninfected cells (20). Since the interruption of viral replication can be achieved by either, the impediment of the viral factory or the death of the virally infected cell, we do not differentiate between synthetic sick and synthetic lethal cellular states.

The feasibility of targeting synthetic lethal partners as a host-based antiviral strategy has been substantiated by two independent studies. Pal, et al. demonstrated that by utilizing existing transcriptomics and CRISPR data, valid synthetic lethal targets can be predicted in the context of SARS-CoV-2 (27). In their study, the authors predicted potential VIHs from a handful of transcriptomics datasets in SARS-CoV-2-infected cell lines and patient tissues. SL partners of these VIHs were then identified from cancer SL pairs based on the ISLE algorithm (28) and prioritized by CRISPR KO screens from SARS-CoV-2-infected cells. The authors validated 24 predicted SL targets demonstrating reduced viral replication in infected cells combined with a decrease in infected cell counts. Similarly, Navare, et al. illustrated that viral-host protein-protein interactions can render host proteins hypomorphic and expose cells to synthetic lethal vulnerabilities (29) Specifically, this study demonstrated that the Golgi-specific brefeldin A-resistance guanine nucleotide exchange factor 1 (GBF1) (30) was converted to a hypomorph by binding to poliovirus 3A protein and that depletion of ADP Ribosylation Factor 1 (ARF1), an SL partner of GBF1, by CRISPR and shRNA led to cell death specifically in 3A-expressing cells, thus reducing viral replication (29). Importantly, GBF1 is a critical proviral factor and a common target of many viruses, suggesting that targeting SL partners of GBF1 could have broad antiviral activity.

Taken together, this evidence suggests that protein-protein interactions and transcriptomic alterations can generate VIHs, the synthetic lethal partners of which could be targeted to inhibit typical viral infection. In this study, we aimed to determine whether various existing omics types can accurately predict VIHs and, by extension, synthetic lethal targets. We also posit that a certain subclass of genes, identified through genome-wide CRISPR KO studies, might contain members that are synthetically lethal with VIHs and thus can be explored as antiviral therapeutics. We expect to find such SL targets among genes depleted in CRIPSR KO screens alongside antiviral genes, since depletion of both of these subclasses of genes should lead to diminished cell viability or cell death.

To explore this, we have engineered a computational pipeline to discern the omics datasets best suited for predicting VIHs and the associated synthetic lethal targets. Our findings indicate that multiple different omics data classes can be used to predict synthetic lethal targets, many of which are incorporated in the pooled survival data of CRISPR KO screens. Furthermore, we highlight that many candidate synthetic lethal targets of SARS-CoV-2 infection are also targets of influenza infection, suggesting that shared synthetic lethal targets may be suitable for host-based therapeutics against multiple viruses.

## MATERIAL AND METHODS

### Data set selection

We utilized 54 published gene sets from 22 studies, encompassing ten different omics data classes (see Table 1 and references under ‘Identification of viral-induced hypomorphs (VIHs)), along with 11 SARS-CoV-2 and three influenza genome-wide CRISPR-Cas9 knock-out screens for this study (see Table 3 and references under ‘Compilation of publicly available CRISPR KO screen data’). We specifically focused on genome-wide screens conducted in cell lines or tissues infected with SARS-CoV-2 or influenza viruses.

**Table 1.**
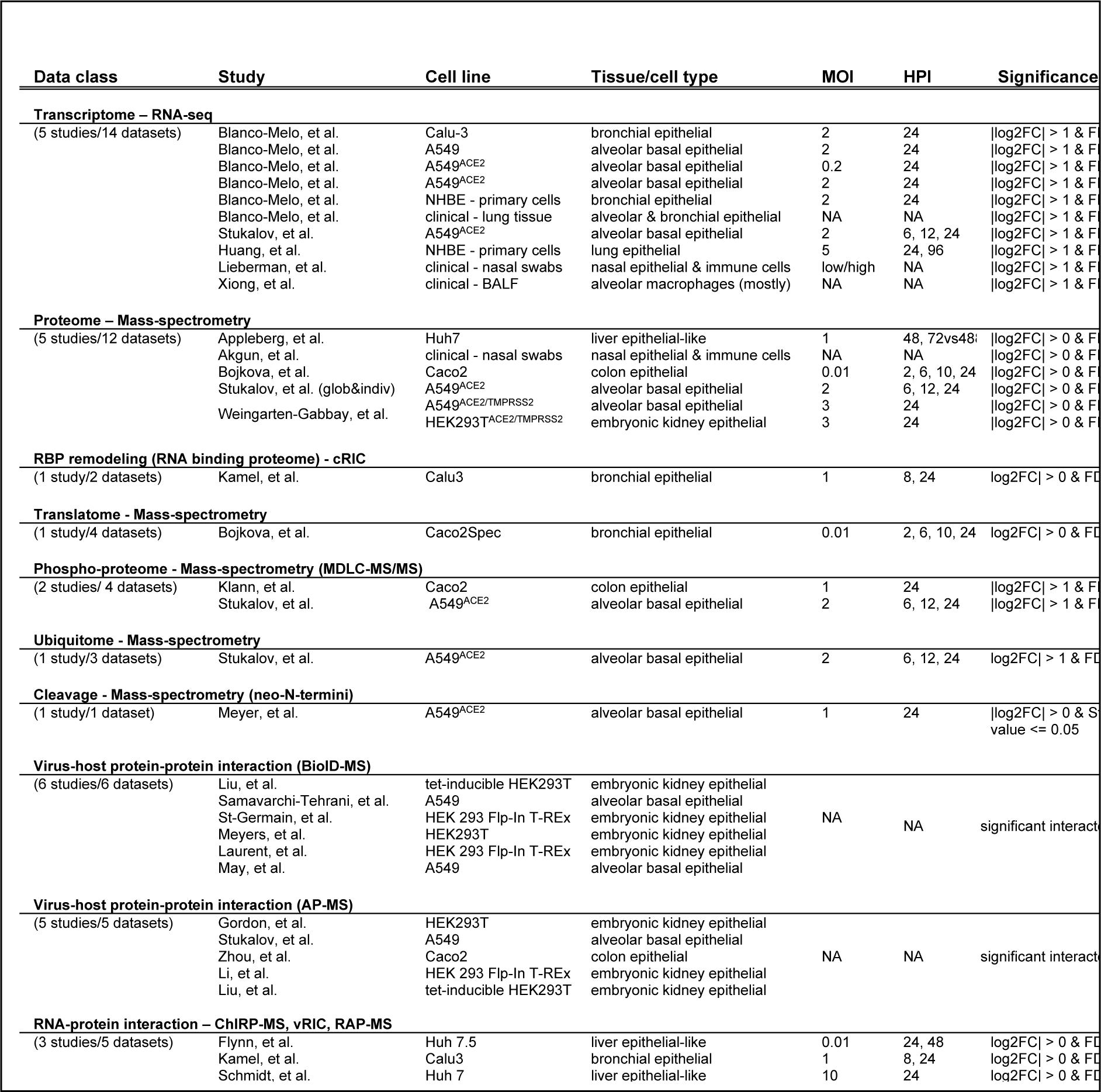
Omics data classes and datasets used to identify candidate synthetic lethal targets based on viral-induced hypomorphs. Table includes experimental methods and tissue/cell type used as well as the multiplicity of infection (MOI) and hours post infection (HPI).

**Table 2.** Number of viral-induced hypomorphs (VIHs) identified among omics data classes, VIHs in SynLethDB, and synthetic lethal (SL) partners. Data classes are ordered by the number of unique SL partners they predicted.

| Data class | N VIHs | N VIHs in SynLethDB | N SL partners of VIHs |
| --- | --- | --- | --- |
| Cleavage | 77 | 60 | 520 |
| Phosphoproteome | 2448 | 1753 | 4491 |
| Proteome (down) | 1244 | 798 | 3463 |
| Proteome (up & down) | 2100 | 1334 | 5048 |
| Proteome (up) | 994 | 621 | 3206 |
| RBP (down) | 107 | 86 | 440 |
| RBP (up & down) | 217 | 175 | 784 |
| RBP (up) | 111 | 90 | 496 |
| Transcriptome (down) | 11263 | 6632 | 9267 |
| Transcriptome (up & down) | 14360 | 7774 | 9484 |
| Transcriptome (up) | 6423 | 3111 | 6280 |
| Translatome (down) | 49 | 26 | 148 |
| Translatome (up & down) | 56 | 31 | 167 |
| Translatome (up) | 7 | 5 | 25 |
| Ubiquitinome | 224 | 176 | 2800 |
| Virus-host PPI (AP) | 2076 | 1419 | 5273 |
| Virus-host PPI (BioID) | 4191 | 2877 | 6184 |
| Virus-host RPI | 312 | 251 | 1175 |
| **Total unique genes** | **15,183** | **8,213** | **9,643** |

**Table 3.**
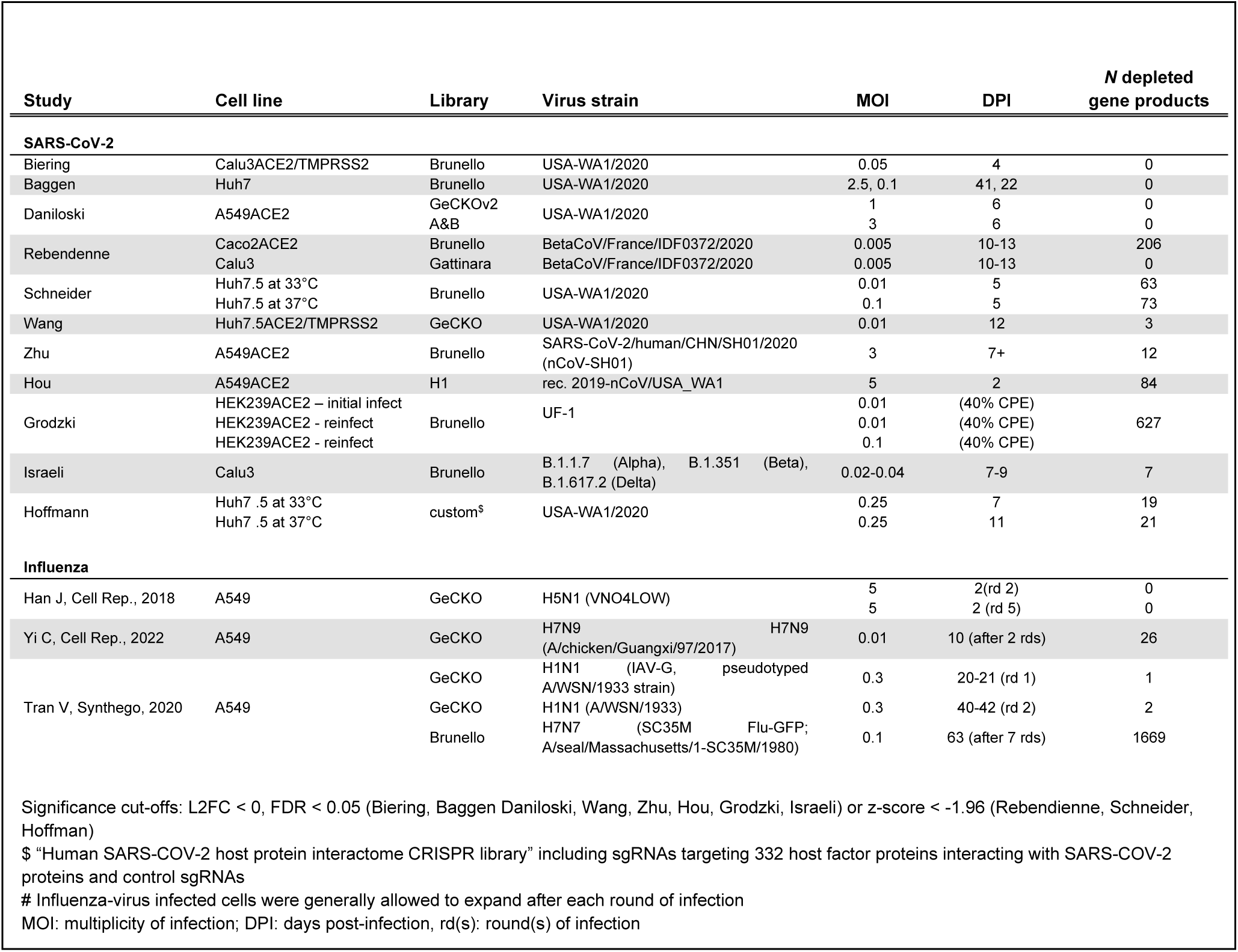
CRISPR KO datasets used for validating predicted synthetic lethal targets.

### Identification of viral-induced hypomorphs (VIHs)

We defined VIHs as gene products that exhibited significant alterations in any of the aspects corresponding to the ten omics data classes when comparing virus-infected cells to non-infected cells. Omics data classes included bulk or single-cell RNA sequencing (transcriptomes) (31–35) specialized mass-spectrometry (proteomes, RNA-binding proteome, phosphoproteomes, ubiquitomes, cleavage) (32, 36–41) as well as combinations of affinity purification (AP-MS)(32, 42–45) or proximity-dependent labeling (BioID-MS) (45–50)with mass-spectrometry (Table 1). We also considered alterations in translation rates (translatome) (40) as well as changes in the host RNA-binding proteome and the host interactome with viral RNAs (51–53). All selected datasets were derived from genome-wide studies in cells infected with live SARS-CoV-2 virus, SARS-CoV-2 pseudovirus, or transduced with individual SARS-CoV-2 virus proteins or RNAs for interactome studies.

When applicable, we categorized VIHs as "up" or "down" based on abundance changes and conducted separate analyses for these subsets in addition to analyzing the complete list of all VIHs. For example, we performed separate directional analyses on transcriptome, proteome, RNA-binding proteome, and translatome data classes for both up- and down-regulated gene products. Whenever available, we adopted the statistical cutoff criteria used in the associated publication of the respective dataset or reported FDR-adjusted *P-*values < 0.05 as significantly altered by infection. Lists of significant interactors for virus-host protein-protein interactions (PPI) and virus-host RNA-protein interactions (RPI) were obtained from the relevant publications. Using these criteria, we identified VIHs for each data class and then determined the union of VIHs across all datasets within each of the ten omics data classes. This union, in addition to directional subsets, were subsequently utilized in our computational pipeline for predicting synthetic lethality targets.

### Prediction of candidate SL targets

We defined candidate SL targets as the SL partners of VIHs listed in SynLethDB 2.0 (54). Thus, we identified all VIHs that also appeared in an SL pair in SynLethDB and consolidated the union of all SL genes associated with those VIHs. This process was performed separately for each list of VIHs compiled from different omics data classes. The resulting list of SL targets was then employed in downstream analyses to evaluate prediction accuracy against depleted products in CRISPR KO studies and to assess prediction agreement among different data classes.

### Compilation of publicly available CRISPR KO screen data

We gathered publicly available data from genome-wide CRISPR KO screens conducted in cells infected with either SARS-CoV-2 (55–65) or influenza virus (66–68) and identified significantly depleted genes. Genes were considered significantly depleted if they exhibited FDR-corrected *P-*values < 0.05 along with negative log_2_ fold-changes or z-scores < -1.96. The aggregated set of CRISPR-depleted genes from all the studies was then used as the surrogate gold standard set of SL targets when evaluating the predictive performance of each omics data class in our study.

### Enrichment analysis of predicted SL genes in CRISPR depleted genes

We used hypergeometric tests to quantify the enrichment of predicted SL targets among depleted genes in CRISPR KO studies. The union of all genes tested across CRISPR KO studies was considered the background population. We considered FDR-adjusted P-values < 0.05, computed using the Benjamini-Hochberg method (69), as indicative of significance.

### Sensitivity, specificity, and precision metrics

To compute sensitivity, specificity, and precision statistics, we designated genes depleted in CRISPR KO screens as positive cases. Negative cases consisted of products evaluated using CRISPR KO screens but were not depleted. Thus, the complete set of genes evaluated by CRISPR KO served as the ground truth against which we assessed SL target predictions. Predicted SL targets generated by our pipeline were considered positive tests, while negative tests consisted of genes not predicted to be SL targets. Therefore, a predicted SL target that was also depleted in CRISPR KO data was considered a true positive. A true negative referred to a gene product not predicted to be an SL target and not depleted in CRISPR KO data.

### Datasets used for characterizing broadly antiviral candidate SL targets

We employed several datasets to further characterize 71 candidate SL target genes that may be broadly antiviral for SARS-CoV-2, influnza A and possibly other viruses. Gene paralogs were obtained from Ensembl using the biomaRt package (70) (version 2.50.3). Extremely multi-functional genes (or hubs) were downloaded from MoonDB 2.0 (71). Essential genes for ten different cell lines were downloaded from supplemental tables in Bertomeu et al. (72). Drugs targeting predicted broadly antiviral SL targets were downloaded from DrugBank (September 22^nd^, 2022) (73).

### Analysis software

All analyses were performed using R (version 4.1.1). Heatmap analyses utilized the pheatmap package (version 1.0.12) with hierarchical clustering based on Euclidean distances. GO:biological process and Reactome enrichment tests were conducted using the clusterProfiler (version 4.2.2) (74) and ReactomePA (75) (version 1.38.0) packages, respectively. For all datasets analyzed in our study, gene and gene product identifiers were aligned with a reference list of gene symbols using the org.Hs.eg.db R package (76).

## RESULTS

### Synthetic lethal target prediction pipeline

Numerous mechanisms exist by which viruses may induce hypomorphic states within host cells that may introduce synthetic lethal (SL) relationships. These include modifications to virus-host protein-protein interactions (PPIs), transcription, protein abundance, post-translational modifications (PTMs), protein repositioning, and proteolytic cleavage, among others (21, 22). Consequently, a myriad of omics-level data acquisition techniques could potentially facilitate the identification of virus-induced hypomorphs (VIHs) and subsequently their SL partners, which could serve as host-based therapeutic targets. To examine this, we developed a computational pipeline to integrate data from various omics data sources, identify potential VIHs, and predict their prospective SL partners as antivirals (Figure 1B). Here, we define VIHs as gene products with impaired function, irrespective of whether the identified molecular species rise or fall in abundance following viral infection, as protein or pathway functionality may be compromised in either scenario.

**Figure 1.**
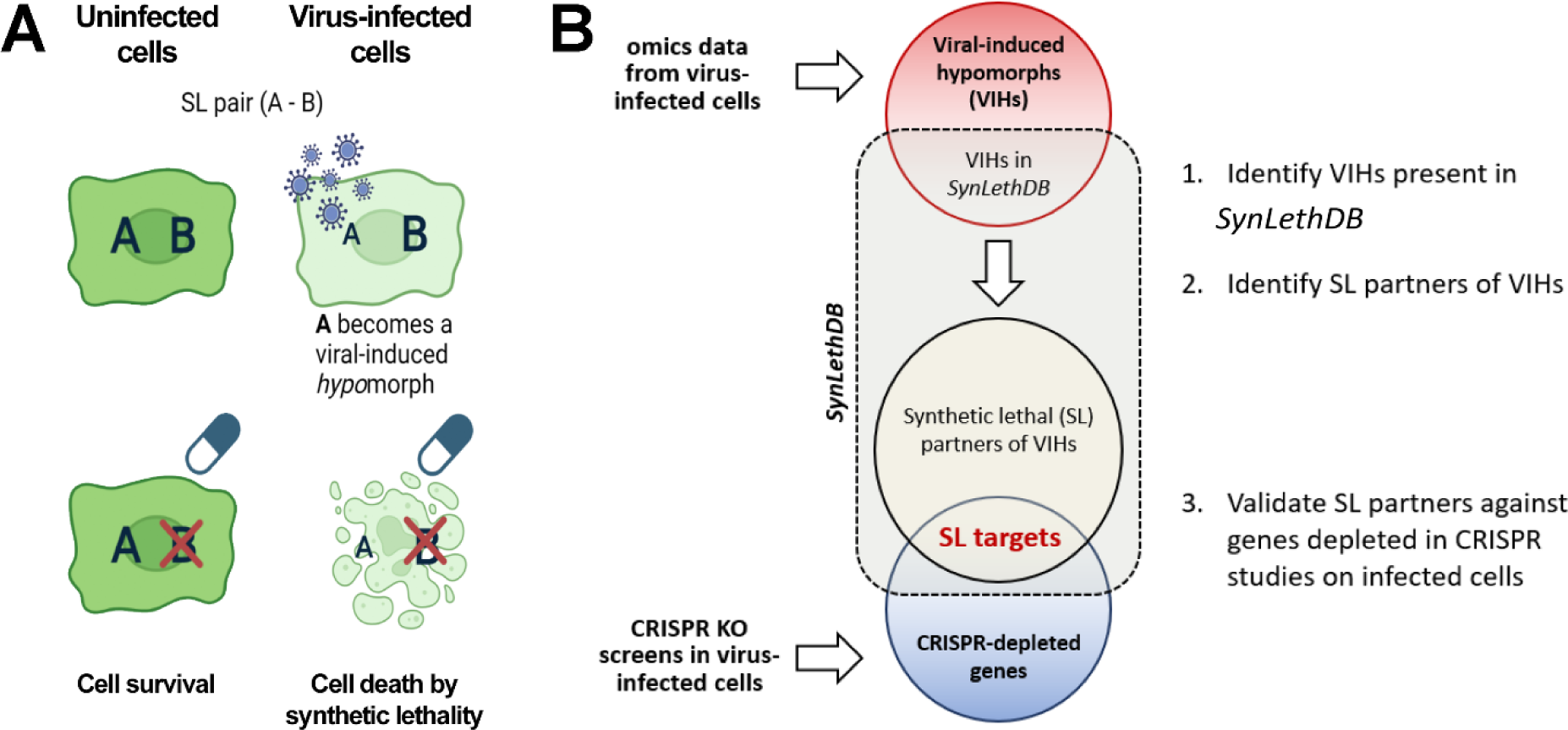
Strategy for identifying antiviral synthetic lethal targets. A) Synthetic lethality in virus-infected cells: In uninfected cells, disrupting one member of an SL pair (A-B) does not impact viability. In infected cells, where one of the SL partners is hypomorphic, disrupting its SL partner can compromise cells, or cause cell death. That is, protein B suppresses the negative consequences of A becoming a viral-induced hypomorph when cells are infected. Conversely, protein A suppresses the negative consequences of drugging B in uninfected cells. However, the combination of a viral-induced hypomorph and targeting its SL partner with a drug has no recourse from these negative consequences and results in lethality. B) Workflow used for validating predicted SL targets from various omics data classes. SL: synthetic lethal; VIH: viral-induced hypomorph.

For each omics data category, we selected individual datasets from as many applicable studies as were available in the literature which led to the inclusion of one to 14 datasets per omics data class (Table 1). We aimed to encompass a large variety of infection conditions (varying cell lines, virus strains, infection multiplicity, and timeline) cited in the literature and thus adopted an aggregate approach incorporating all available datasets within each data class. Following this, we applied a filtering methodology to identify the candidate SL genes with the highest likelihood of successful experimental validation (Figure 1).

Initial identification of putative VIHs was achieved by finding molecular species (transcripts, protein abundance, or PTMs) that exhibited significant alterations following SARS-CoV-2 infection. Subsequently, the SL partners of these VIHs were identified within the SynLethDB resource (54), a public repository of SL pairings developed for cancer research. Since genome-wide CRIPSR KO screens in the setting of viral infection should contain SL genes due to reduced viability of cells with SL gene knockouts, we used depleted CRISPR KO gene pools to quantify the power of each omics data class for predicting SL targets. These measures included enrichment of candidate SL targets as well as specificity, sensitivity, and precision statistics. Finally, we used the same depleted CRISPR KO gene pool as a filter to further prioritize candidate SL targets. This step also filtered out the genes in the CRIPSR KO-depleted gene set that might not be in SL relationships such as antiviral genes and genes that are essential for cell growth and survival.

The remaining proteins represent candidate SL targets that have led to decreased cell viability when knocked-out during viral infection compared to infection alone in CRISPR-based KO screens in at least one published viral infection scenario.

### Data selection

We curated datasets from published research focusing on SARS-CoV-2 and influenza viruses due to their ongoing global health and pandemic implications. Given the intensive research on these viruses and their relatives, a considerable number of datasets describing various aspects of gene functionality for orthomyxo- and coronaviruses are available, which is ideal for an exhaustive exploration of the most accurate predictors for virus-specific SL targets from a practical perspective. In total, we compiled 56 omics datasets from 22 distinct studies on SARS-CoV-2 infection, which analyzed a wide spectrum of cellular changes (Table 1). The datasets depict host interactors of viral proteins and RNA (32, 42, 44–53, 77) infection-induced alterations in host gene expression (31–33, 35, 78) and protein abundance (32, 36–38, 40), changes in translation rates (40), as well as alterations in post-translational modifications (32, 39), cleavage (41), and the RNA-binding proteome (RBP) (51), comprising ten VIH data classes in total. Host interactors of viral proteins were further split based on technology: affinity purification (AP) or a proximity-based approach with tagged target proteins (BioID), both combined with mass spectrometry (MS) (Table 1). Both techniques used individual viral open reading frames (ORFs) as bait to identify interacting host proteins. However, published BioID-MS data were derived from cells both with and without viral infection. AP-MS was predominantly employed to capture virus-host multi-protein complexes (79), whereas BioID-MS (80) serves as a complementary method to examine transient interactions based on physical proximity, a frequent occurrence throughout the viral infection cycle (81).

### Synthetic lethal target prediction

#### Identification of VIHs and their SL partners from omics data stemming from virus-infected cells

Our pipeline’s initial step (Figure 1B) involved the identification of potential VIHs by selecting gene products significantly impacted by viral infection, as evident from the ten omics data classes outlined in Table 1. For VIH identification, we relied on the significance criteria stipulated in each dataset’s corresponding publication. We sub-categorized VIHs based on the increase or decrease in molecular species abundance post-infection for transcriptomics, proteomics, RNA-binding proteomics, and translatomics data. For phosphoproteome, ubiquitome, and protein cleavage data, we ascertained the total absolute change for each protein based on the aggregate changes reported for all pertinent peptides.

Following VIH identification from each data class, we narrowed down the VIHs to those also present in the commonly utilized SynLethDB version 2.0, and then identified the SL partners of putative VIHs (Figure 1B, Table 2). SynLethDB (54) comprises a collection of SL pairs procured from diverse sources, including confirmed SL pairs from the literature based on genome-wide siRNA, CRISPR KO screens, or bi-specific SL shRNA screens, in addition to predictions from computational algorithms, text mining, and SL pairs from other similar databases. The SynLethDB version utilized in this study contains 35,943 human SL pairs, encompassing 9,855 unique SL genes. On average, each gene is in a SL relationship with three other genes.

Overall, across all data classes, we identified a total of 15,183 unique VIHs and a total of 9,643 unique SL partners for those VIHs (Table 2). The number of unique VIHs and SL partner targets identified using each data class differed substantially (Table 2, Figure 2). The maximum numbers of VIHs and SL partners were derived from transcriptomic, PPI, and proteomic studies, while the minimum were derived from translatome and cleavage studies. We found significant correlations between the number of datasets in a class and the total number of predicted VIHs (Pearson coefficient = 0.80, P-value = 0.006) as well as their predicted SL partners (Pearson coefficient = 0.83, P-value = 0.003). This provides evidence, as anticipated, of the poor overlap between individual datasets within each data class, resulting from the broad spectrum of different infection scenarios we included. The higher numbers of predicted VIHs stem from the fact that each additional dataset adds new and different altered genes to the overall VIH gene pool when datasets don’t agree very well. We chose this strategy because it provided us with a suitably comprehensive list of candidate SL targets from a larger variation of potentially relevant infection scenarios for further evaluation.

**Figure 2.**
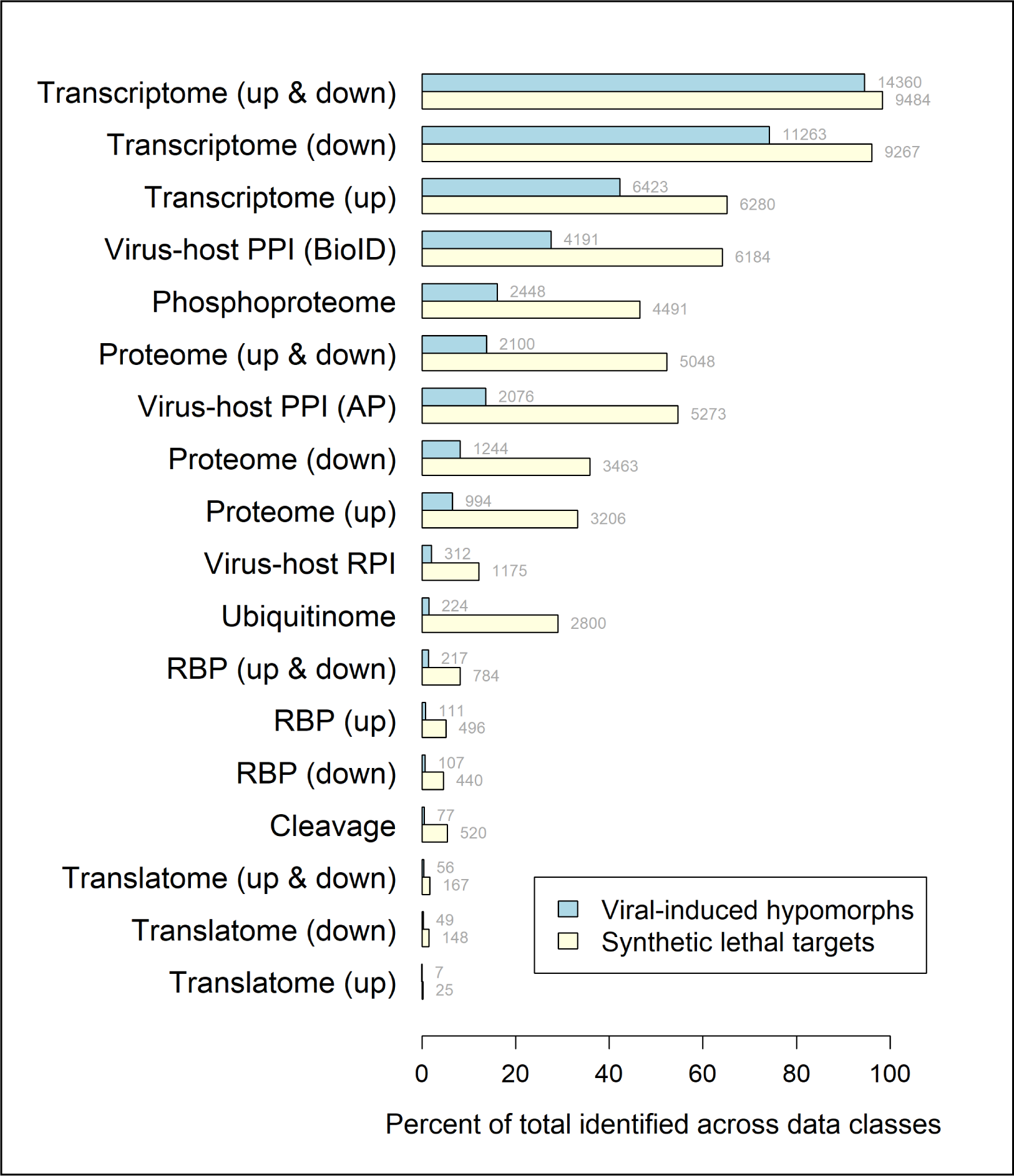
Percentage of predicted viral-induced hypomorphs and synthetic lethal targets for individual omics data classes. For each class, bar width indicates the percentage of VIHs (blue) and SL targets (yellow) predicted out of the combined set of all predictions across data classes. For directional omics data, results are shown for gene products that increased (“up”) or decreased (“down”) in abundance. The numbers at the ends of the bars indicate absolute number of VIHs and SL targets identified.

#### VIHs and SL targets inferred by transcriptomics encompass nearly all genes predicted across omics data classes

To ensure our synthetic lethality prediction pipeline did not overlook valid SL targets, we examined whether consolidating predictions from multiple omics data classes would identify more valid targets compared to employing any single data class. In other words, we explored whether the combined usage of multiple omics data classes would result in a more sensitive prediction of valid SL targets, given that different classes probe different facets of infection and might yield non-overlapping predictions. However, our findings showed that 95% of all the VIHs predicted across data classes were included in the predictions derived from transcriptome data, and 98% of all the SL targets predicted across classes were covered (Figure 2). Therefore, within the scope of the data classes we explored, transcriptome data provides a near-maximal level of sensitivity in predicting SL targets. The downside is that the transcriptome data class, as elaborated below, demonstrated the lowest statistical precision, i.e., the highest proportion of false-positive predictions, among the data classes (Table 4).

**Table 4.** Metrics of performance among omics data classes for predicting valid synthetic lethal targets against SARS-CoV-2 infection. The total number of CRISPR-depleted non-redundant candidate SL targets is 738.

| Data class | N candidate SL targets<br>also CRISPR-depleted | Enrichment of SL targets in<br>CRISPR-depleted genes | Sensitivity | Specificity | Precision |
| --- | --- | --- | --- | --- | --- |
| | | $-\log_{10}(\text{FDR-adjusted } P\text{-value})$ | | | |
| Virus-host PPI (BioID) | 600 | 85.79 | 0.58 | 0.72 | 0.1 |
| Virus-host PPI (AP) | 526 | 74.7 | 0.51 | 0.76 | 0.1 |
| Transcriptome (down) | 732 | 71.69 | 0.71 | 0.58 | 0.08 |
| Transcriptome (up & down) | 736 | 68.6 | 0.71 | 0.57 | 0.08 |
| Proteome (up & down) | 490 | 63.63 | 0.47 | 0.77 | 0.1 |
| Transcriptome (up) | 555 | 60.68 | 0.54 | 0.72 | 0.09 |
| Proteome (down) | 378 | 58.08 | 0.37 | 0.85 | 0.11 |
| Virus-host RPI | 188 | 48.94 | 0.18 | 0.95 | 0.16 |
| Phosphoproteome | 418 | 46.26 | 0.41 | 0.8 | 0.09 |
| Proteome (up) | 330 | 43.45 | 0.32 | 0.86 | 0.1 |
| RBP (up & down) | 125 | 31.78 | 0.12 | 0.97 | 0.16 |
| RBP (down) | 90 | 31.02 | 0.09 | 0.98 | 0.2 |
| Ubiquitinome | 268 | 28.68 | 0.26 | 0.87 | 0.1 |
| Cleavage | 93 | 27.41 | 0.09 | 0.98 | 0.18 |
| RBP (up) | 74 | 17.15 | 0.07 | 0.98 | 0.15 |
| Translatome (up & down) | 17 | 2.43 | 0.02 | 0.99 | 0.1 |
| Translatome (down) | 15 | 2.21 | 0.01 | 0.99 | 0.1 |
| Translatome (up) | 4 | 1.5 | 0 | 1 | 0.16 |

We anticipated that the directional changes documented in omics datasets (e.g., upregulated versus downregulated transcription) would identify mutually exclusive VIHs. While this was predominantly the case, a considerable number of gene products were predicted to both increase and decrease in abundance. This can be attributed to the fact that a gene product might be overexpressed in one dataset but downregulated in another within the same data class depending, for example, on the viral effect on a particular cell line used in a specific experiment.

#### Convergence of different omics data classes on predicted SL targets despite low agreement for VIHs

Different omics data classes under investigation represent unique perspectives of a virus’s impact on cellular processes, with an uncertainty surrounding the level of agreement across molecular species impacted by varied mechanisms in response to viral infection. Our study examined the overlap of predicted VIHs and synthetic lethality targets across these different omics data classes. By iteratively pairing each data class with others, we calculated the Jaccard index for predicted VIHs. Although the alignment between data classes for predicted VIHs was typically low, a significantly higher degree of congruence was observed for predicted SL targets derived from VIHs (Figure 3). This indicates that despite disparate VIH predictions, subsets of data classes still converged on common sets of SL targets. Our analysis revealed two primary clusters of data classes showing high agreement in SL predictions, with the first encompassing ubiquitinome, phosphoproteome, transcriptome, proteome, and virus-host protein-protein interaction (PPI) classes, while the second included the translatome, cleavage, virus-host RNA-protein interaction (RPI) and RNA binding protein (RBP) classes. Within this second cluster, a high similarity was observed between virus-host RPI and RBP classes, both based on RNA binding. Not surprisingly, the closest agreement for SL partners existed between virus-host PPIs derived from AP and BioID in the first cluster.

**Figure 3.**
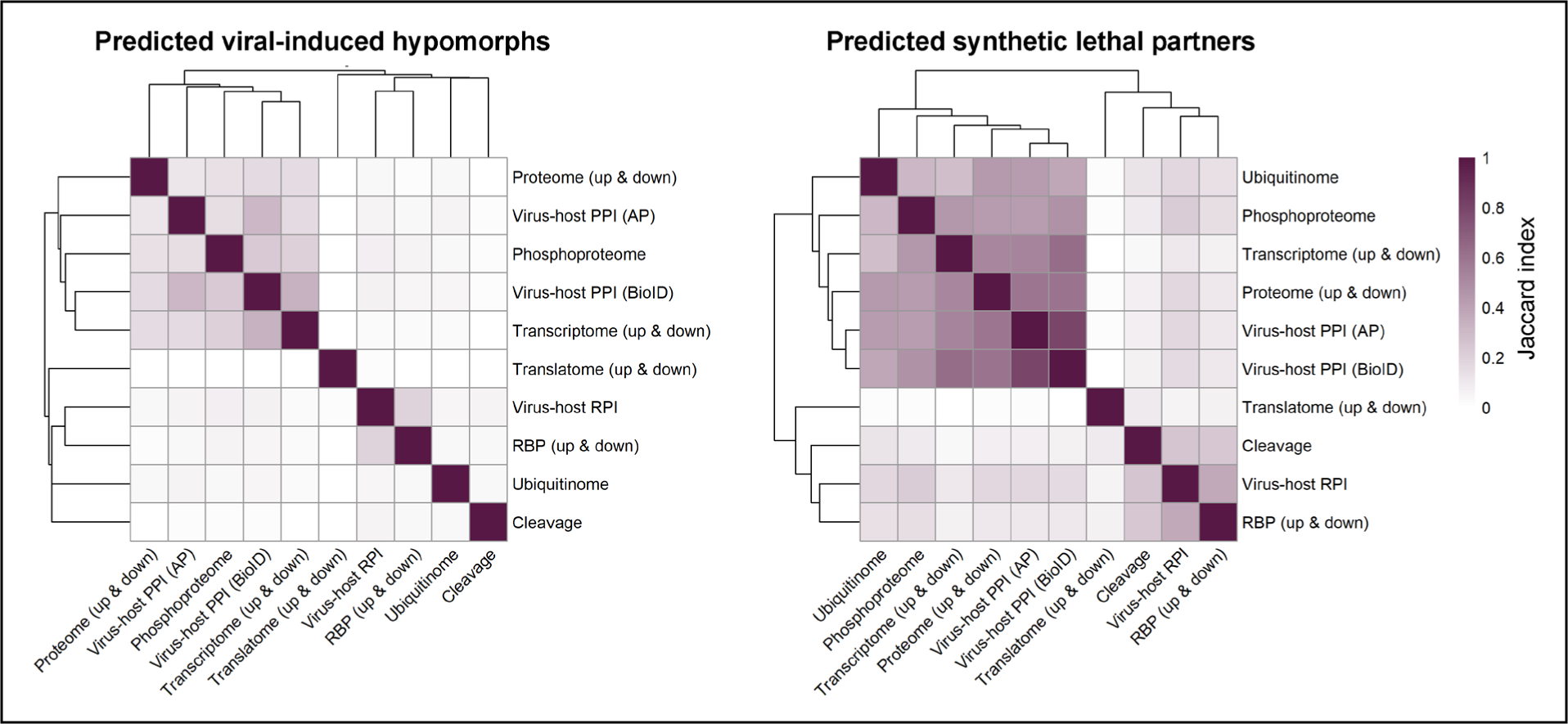
Heatmaps showing pairwise overlap of predicted viral-induced hypomorphs and SL partners across omics data classes. For each data class pairing, the degree of overlap of predicted hypomorphs (left) and synthetic lethal partners (right) was quantified using the Jaccard index with darker purple indicating more significant overlap. The dendrograms on both axes indicate similar groupings of overlapping genes between omics data classes.

### Performance of Omics Data Classes in Predicting Valid Synthetic Lethal Targets

A central aim of our study was to evaluate the predictive accuracy of each data class in identifying legitimate, experimentally verified SL targets. Unfortunately, a universally accepted gold standard for validating SL targets is currently unavailable. To create a representative pool of SL targets, that could serve as a surrogate gold standard bolstered by adequate experimental evidence, we aggregated depleted genes from multiple, diverse CRISPR knockout (KO) screens. However, we note the unknown percentage of false negatives such as antiviral and potentially some essential genes associated with these data. Moreover, there were no genes that exhibited universal depletion across all CRISPR KO datasets, potentially due to variable cell type selection among other experimental variables. As with our omics datasets, we opted to pool depleted genes from multiple disparate CRISPR screens to ensure representation of diverse SARS-CoV-2 infection scenarios. With these caveats, we used the list to assess the performance of each data class for predicting SL targets in SARS-CoV-2 infection. Performance measures included hypergeometric enrichment tests as well as sensitivity, specificity, and precision metrics.

Our analysis confirmed enrichment of SL targets among genes depleted in CRISPR KO studies, regardless of the employed omics data class (Table 4). This validates prior research, demonstrating that genes depleted in CRISPR KO studies indeed encompass authentic SL targets which, upon disruption, selectively diminish the viability of infected cells (27). We observed that SL targets predicted using BioID-MS virus-host PPI data showed the highest enrichment in CRISPR KO-depleted genes. When directional changes were considered, enrichment scores among SL targets derived from sets of gene products that were decreased in abundance outperformed those with increased levels. However, both RNA and protein abundance changes in either direction resulted in highly significant CRISPR KO-depletion enrichment, suggesting that either could create VIHs in an infected cell, thereby sensitizing the cell to SL disruption.

Sensitivity and specificity scores spanned a broad range across data classes, with the highest sensitivity observed in the transcriptome, virus-host PPI and proteome classes and the highest specificity within the translatome, RNA-binding proteome and cleavage data classes (Table 4, Supplementary Figure S1). However, the statistical precision of all data classes for the prediction of SL targets was less than or equal to 0.20 (Table 4), indicating that only a proportion of predicted SL partner targets were valid. We hypothesized that limiting predicted SL targets to those identified across multiple data classes might improve the statistical precision of our predictions without compromising sensitivity. However, we found that the maximum precision achievable with this approach was 0.22 and required SL targets to be predicted in 90% of data classes, at the cost of reducing prediction sensitivity to 0.05. Overall, this implies that predicting SL targets from any omics data class investigated here requires further filtering of candidate SL targets through data from studies validating the impact of target disruption on cell viability (such as CRISPR KO and/or siRNA data), as a critical step of our pipeline.

We also probed whether the number of data sets compiled for each omics data class correlated with the different metrics collected using the ‘gold standard’ CRISPR-depleted gene set. Although this correlation was not significant for hypergeometric enrichment scores (Pearson coefficient = 0.55, P-value=0.10), it significantly correlated with sensitivity (Pearson coefficient = 0.79, P-value = 0.007), specificity (Pearson coefficient = -0.83, P-value = 0.003), and precision (Pearson coefficient = -0.67, P-value = 0.03).

Upon examining the pattern of positive predictions for SL candidate genes that were also CRISPR KO-depleted SL targets, we identified two distinct data class clusters (Figure 4). These clusters largely reflected the groupings observed in our Jaccard index analysis (Figure 3), further illustrating a convergence in predicted SL targets. The two primary clusters share overlapping SL targets but differ in the number of SL targets predicted. Overall, the cluster consisting of transcriptome, virus-host PPI, phosphoproteome, proteome, and ubiquitinome classes predicted a greater number of valid SL targets than the second cluster consisting of virus-host RPI, RBP, cleavage, and translatome data classes. While this second cluster identified a smaller number of candidate SL targets (lower sensitivity), its members predict a higher proportion of true positives (higher precision) (Table 4).

**Figure 4.**
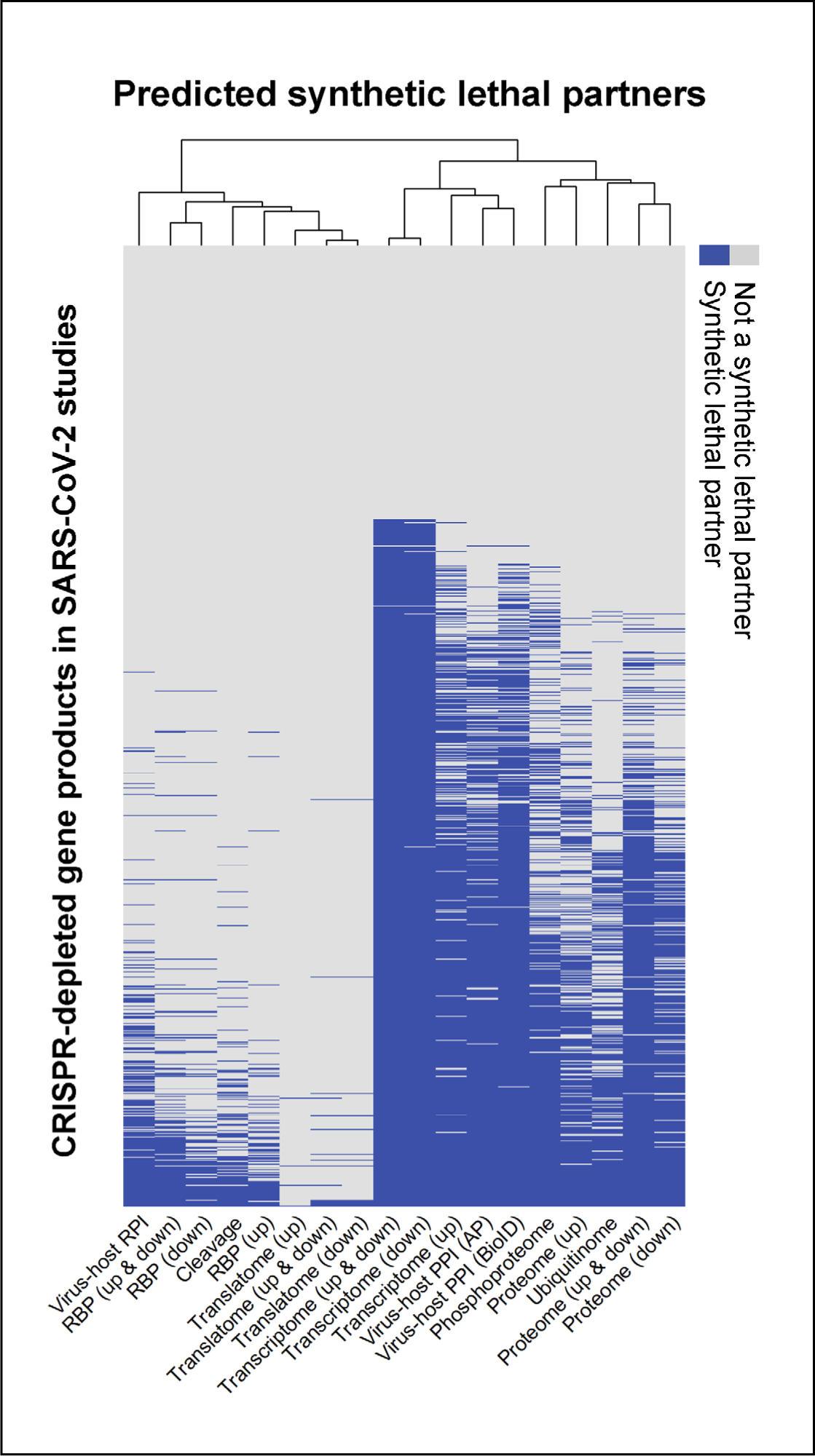
Candidate synthetic lethal targets across omics data classes depleted in SARS-CoV-2 CRISRP KO studies. Binary heatmap showing which of the 1,032 genes in the SARS-CoV-2 CRISPR-depleted gene pool (rows) were predicted to be SL targets for each omics data class. Predicted targets are indicated in blue (refer to Supplementary Tables S6 and S7 for gene names). Grey cells indicateCRISPR-depleted gene products not predicted to be an SL target. Rows are sorted by number of omics data classes predicting the product as an SL target.

Our results highlight a substantial proportion of CRISPR-depleted genes that were not predicted to be SL partners by any omics data class (28%; N=294; Tables 2 and 4, Figure 4), primarily due to their absence in SynLethDB. These genes might hold undiscovered SL relationships with VIHs in the context of SARS-CoV-2 infection or might represent antiviral genes whose disruption enhances viral propagation. The remaining 738 genes are higher-confidence SL targets in the context of viral infection, substantiated by three lines of evidence forming part of our pipeline: 1) experimental evidence of depletion in a CRISPR screen of virus-infected cells, indicating that their disruption adversely affects cell survival specifically in virus-infected cells; 2) evidence of being in a negative genetic relationship with a gene shown to undergo significant alteration due to viral infection; and 3) evidence of being part of a SL pair in the alternate context of altered cellular states due to cancer.

### Functional profiling of high-confidence SL targets

We performed GO:biological process and Reactome enrichment analyses to identify the cellular pathways and functions associated with the 738 high-confidence SL targets mentioned above. There were 264 GO:biological process classes enriching for the SL targets, and they were primarily associated with the cell cycle, nucleic acid processing (including RNA splicing/processing and many rRNA-related processes), metabolism, chromosome organization, intracellular localization/transport as well as organelle biogenesis (Supplementary Table S1). Reactome pathways showing enrichment (N=209) were primarily associated with mitotic processes, DNA repair, RNA splicing/processing, translation, infectious disease processes, RNA polymerase II transcription, signal transduction, autophagy, transport mechanisms, organelle biogenesis, metabolism, and stress responses (Supplementary Table S2).

Overall, these findings align well with those of Pal et al. in associating cellular stress responses, RNA splicing/processing, DNA repair, and translation with SL targets (27). However, because our results are based on larger and more diverse datasets, they revealed a broader array of biological functions associated with SL targets in the context of SARS-CoV-2 infection and, importantly, included multiple GO terms and pathways corresponding to viral infection themes that included the entire viral life cycle.

### Identification of common SL targets for SARS-CoV-2 and Influenza viruses

If pan-viral SL targets exist, we would expect that some of the synthetic lethal targets we predicted using SARS-CoV-2 omics data, would also be present in CRISPR-depleted genes derived from cells infected with other viruses. We chose influenza viruses because they are another global viral threat and also quite different from coronaviruses. For example, coronaviruses have a continuous dsRNA genome, bind to ACE2 for cell entry and conduct replication and transcription in the cytoplasm (82). Influenza viruses use sialyloligosaccharides as their main entry receptors, have a unique segmented genome, and replicate in the host cell nucleus (83). Hence, we assembled a pool of 1,693 unique genes that were significantly depleted in three influenza CRISPR KO studies and determined whether our SARS-CoV-2 SL targets were significantly enriched among these genes (Table 5). We found that, although enrichment scores were substantially lower than for SARS-CoV-2 CRISPR-depleted genes, half the data classes showed significant scores (transcriptome, virus-host PPI, proteome and phospho-proteome). Similar to SARS-CoV-2 predictions, precision based on influenza CRISPR KO depleted genes was uniformly low. Sensitivity and specificity values indicated that predictions were not substantially more accurate than would be expected by chance.

**Table 5.** Metrics of performance for predicting valid synthetic lethal targets against influenza infection using omics data from SARS-CoV-2 infections. The total number of non-redundant CRISPR-depleted candidate SL targets is 892.

| Data class | Enrichment of SL targets in<br>CRISPR-depleted genes | Sensitivity | Specificity | Precision |
| --- | --- | --- | --- | --- |
| | $-\log_{10}(\text{FDR-adjusted } P\text{-value})$ | | | |
| Transcriptome (down) | 3.07 | 0.51 | 0.54 | 0.09 |
| Transcriptome (up & down) | 3.07 | 0.52 | 0.53 | 0.09 |
| Virus-host PPI (BioID) | 2.22 | 0.34 | 0.69 | 0.09 |
| Proteome (up & down) | 1.9 | 0.28 | 0.75 | 0.09 |
| Proteome (up) | 1.83 | 0.18 | 0.84 | 0.1 |
| Transcriptome (up) | 1.83 | 0.34 | 0.69 | 0.09 |
| Proteome (down) | 1.43 | 0.19 | 0.83 | 0.09 |
| Virus-host PPI (AP) | 1.43 | 0.29 | 0.74 | 0.09 |
| Phosphoproteome | 1.34 | 0.24 | 0.78 | 0.09 |
| Ubiquitinome | 0.9 | 0.15 | 0.86 | 0.09 |
| RBP (up & down) | 0.81 | 0.05 | 0.96 | 0.1 |
| RBP (up) | 0.77 | 0.03 | 0.98 | 0.1 |
| RBP (down) | 0.71 | 0.03 | 0.98 | 0.1 |
| Cleavage | 0.63 | 0.03 | 0.97 | 0.1 |
| Virus-host RPI | 0.63 | 0.06 | 0.94 | 0.09 |
| Translatome (up) | 0.14 | 0 | 1 | 0.08 |
| Translatome (down) | 0.09 | 0.01 | 0.99 | 0.07 |
| Translatome (up & down) | 0.09 | 0.01 | 0.99 | 0.07 |

However, fifty-three percent of the influenza CRISPR KO-depleted genes were predicted as SL targets by one or more data classes utilizing the SARS-CoV-2 omics data-based predictions (Figure 5). Given this substantial overlap as well as the significant enrichment of candidate SARS-CoV-2 SL targets for half the omics data classes in influenza CRISPR KO data, we reasoned that there may be a group of SL targets common to both viral infections. We therefore examined the overlap between the significantly depleted genes in both the SARS-CoV-2 and influenza CRISPR KO data and found 99 shared genes of which 71 (72%) were predicted to be SL targets. These 71 genes may therefore represent broad viral SL targets. GO:biological process terms enriched for these 71 genes included ribosomal biogenesis, ncRNA processing (including rRNA), translation, regulation of sister chromatid cohesion, and DNA replication (Supplementary Table S3). Enriched Reactome pathways largely fell within the same categories but also included influenza/HIV infection, starvation responses, amino acid/protein metabolism, DNA synthesis/repair, nonsense-mediated decay, transcription, mRNA splicing/processing, and TP53-mediated transcriptional regulation. (Supplementary Table S4).

**Figure 5.**
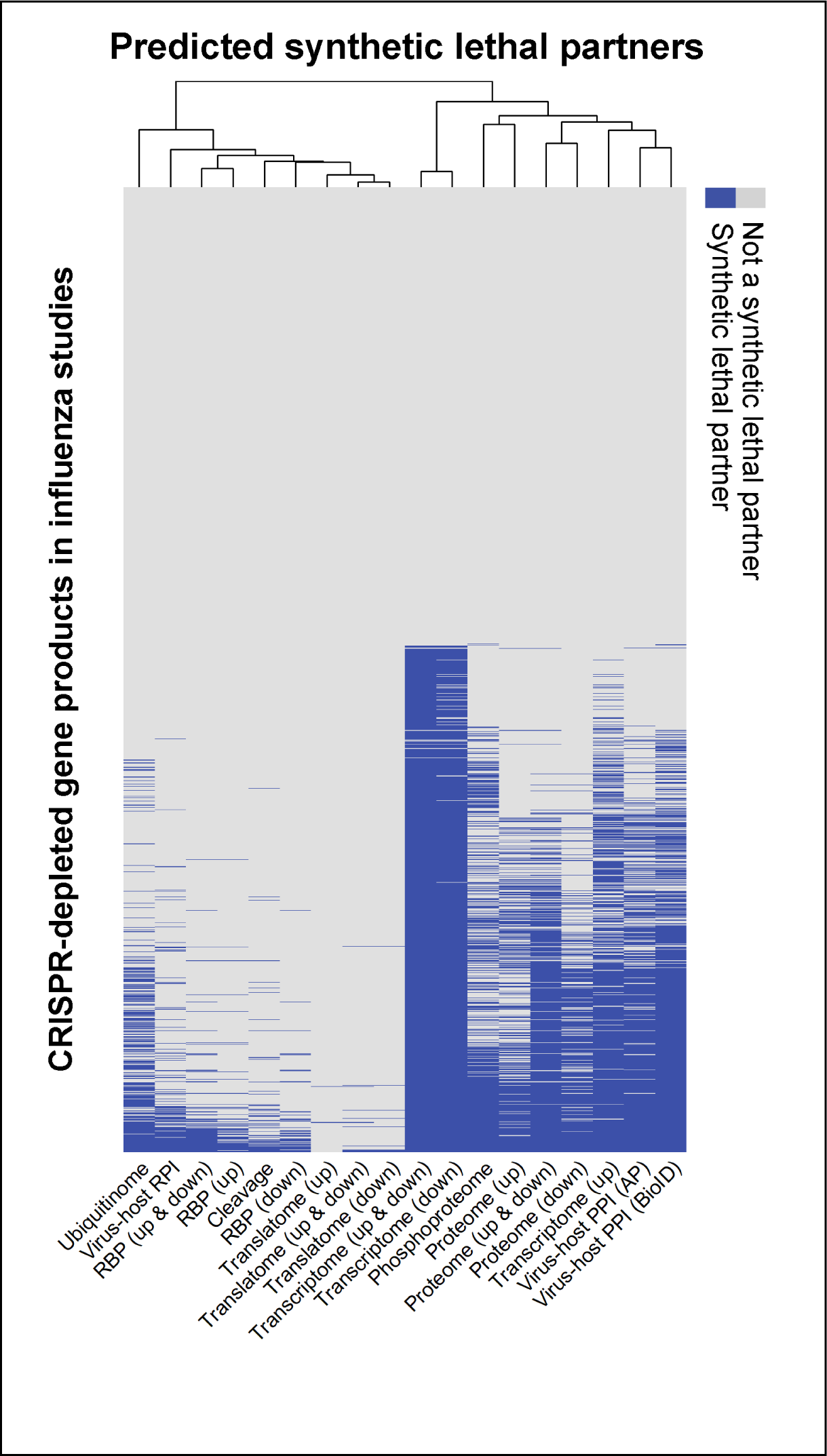
Candidate synthetic lethal targets across omics data classes depleted in Influenza A CRISRP KO studies. Binary heatmap showing which of the 1,693 genes in the influenza CRISPR-depleted gene pool (rows) were predicted to be SL targets for each omics data class. As in Figure 4, predicted targets are indicated in blue (refer to Supplementary Tables S8 and S9 for gene names). Grey cells indicate CRISPR-depleted products not predicted to be an SL target. Rows are sorted by number of data classes predicting the product as an SL target.

Three of the broadly antiviral candidates were also predicted by Pal, et al. as SL targets for SARS-CoV-2 including RPP25 (Ribonuclease P And MRP Subunit P25), a protein involved in tRNA and rRNA processing as well as mRNA metabolism, PCBp1 (Poly(RC) Binding Protein 1), which controls certain steps of pre-mRNA processing, translation, and ferroptosis as well as the initiation of viral replication and viral translation, and MTBP (MDM2 Binding Protein) a protein involved in the initiation of DNA replication, mitotic progression and chromosome segregation, as well as in cell migration and the suppression of invasive behavior due to its interaction with MDM2.

The 71 broadly antiviral candidate SL targets need to be further scrutinized and prioritized before undergoing experimental testing. For example, researchers may wish to deprioritize targets that have high functional interconnectivity risking off-target consequences, such as those referenced in the MoonDB database (71). Other candidate targets that might better be avoided are genes that have documented essentiality in a specific cell line of interest or in multiple cell lines, because their disruption would be expected to kill both infected and uninfected cells (72). Candidate SL genes with numerous paralogs may also warrant careful consideration; if paralogs provide too much functional redundancy, they might prevent efficient synthetic lethality-based killing. However, pairs of genes that are paralogs of each other have been demonstrated to be in SL relationships more frequently than by chance (84). VIH-SL pairs may be prioritized based on any or all these criteria and final candidate SL genes may be chosen for experimental testing based on druggability and availability of drugs or ligands (Table 6).

**Table 6.**
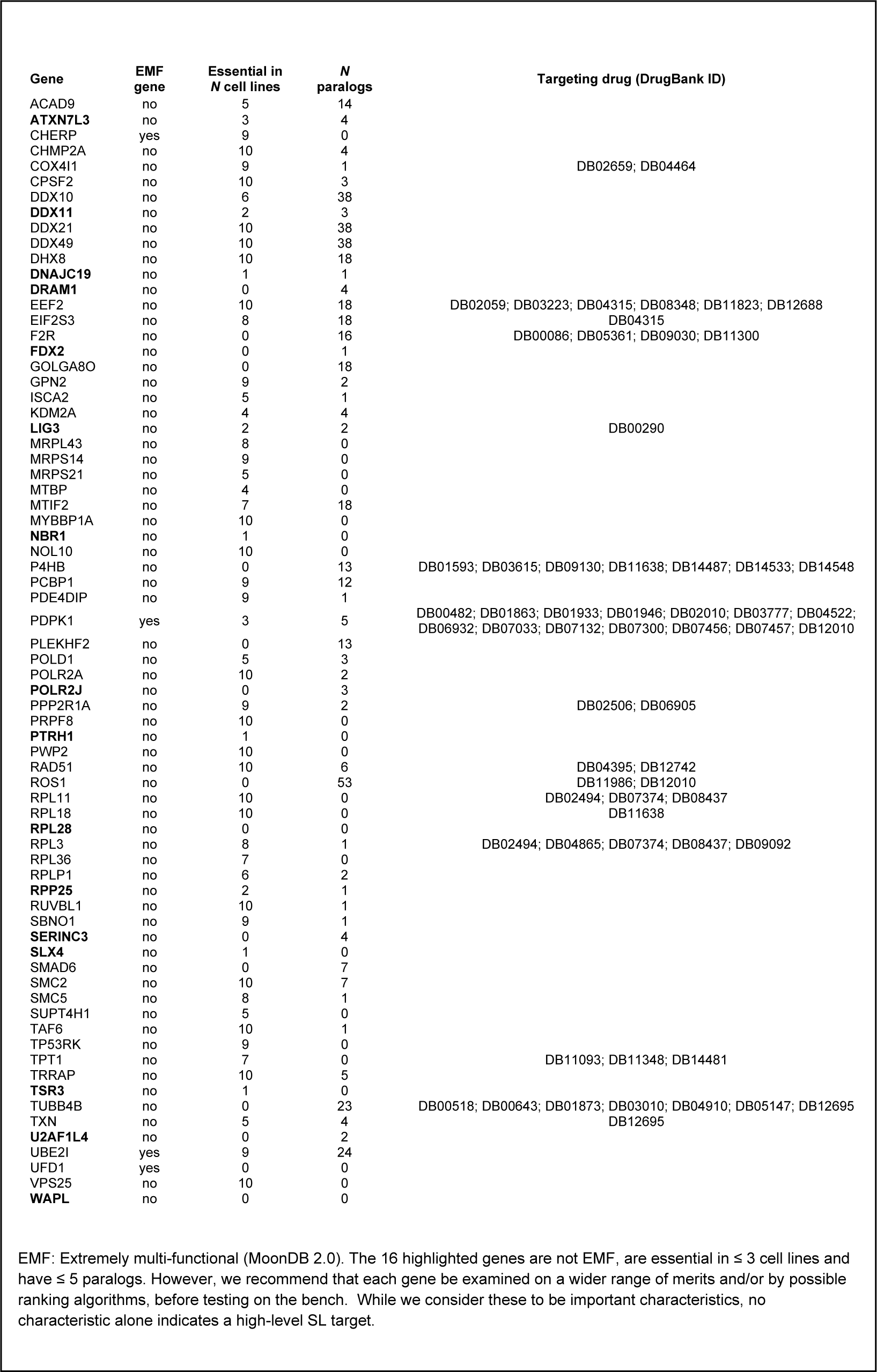
Characteristics of candidate pan-viral synthetic lethal targets potentially useful for prioritizing their confirmation.

## DISCUSSION

This study is part of an on-going effort to examine the concept of synthetic lethality as a means to discover novel host-based antiviral therapeutics. A critical step in this effort is the establishment of a computational pipeline for predicting SL targets. Since synthetic lethality is a type of genetic interaction relatively unexplored in infectious diseases, such a pipeline, once fully established, would expand the number of antiviral drug targets as well as add to the knowledge base of pathogen biology. Our goal is that our pipeline will ultimately generate a plethora of SL targets to be examined, prioritized, and experimentally validated. This study addresses several critical research issues that arose during the process.

Our pipeline generated a list of potential SL targets that are enriched with genes depleted in CRISPR KO studies specific to SARS-CoV-2 infection. These targets were predicted by reusing an array of available omics datasets, which were specific for virus-infected cells, in combination with a synthetic lethality database. One assumption we made was that the SL partners of genes undergoing significant changes in virus-infected cells would be depleted in CRISPR KO screens, thus potentially serving as SL targets. Our results supported this assumption. An approach that would further aid in the prediction of SL targets from CRISPR KO screens is the addition of viral replication measurements in addition to cell viability, a practice not typically adopted for genome-wide functional screens.

We also investigated whether different classes of omics data would differ in their ability to accurately predict SL targets by measuring whether SL prediction by some data classes was better supported by CRISPR KO screens than others. Exploring various omics classes, each interrogating different facets of host-virus biology, we observed that despite varying VIH predictions, there was substantial overlap in the SL vulnerabilites predicted from different omics classes. This also indicates that a common set of biological pathways and functions may be altered upon SARS-CoV-2 infection. Our findings demonstrate that valid SL targets can indeed be derived from different omics data types. Furthermore, we speculate that omics datasets describing other ways in which viral infection alters normal gene and protein functions, such as changes in RNA splicing or additional post-translational modifications, might also be beneficial for SL prediction, but they are currently fewer in number.

Analyzing the high-confidence SL targets predicted by different omics classes, we identified two primary clusters. The larger cluster included classes associated with transcription and translation, post-translational modifications, and host-viral protein interactions. The second cluster included classes associated with translation rate, cleavage, and RNA-binding behavior. We found that each data class varied in sensitivity and specificity in predicting SL targets based on functional assays such as CRIPSR KO screens (Figure S1), indicating that some omics data classes might be more suitable than others for accurate SL prediction. The fact that overall statistical precision was low across all data classes (Table 4), also emphasizes the need to reduce false positives. One way to do so is to limit predicted SL targets to those found depleted in CRISPR KO studies.

Recent findings by Pal et al. (27) confirmed that transcriptomic data alone could be used to predict SL target genes leading to reduced viral replication and cell death in SARS-CoV-2-infected cells when depleted. However, the authors reported a relatively modest difference in cell viability between infected and uninfected cells for at least half of their 26 top candidates. We noted substantial overlap in enriched cellular pathways and functions between our study and theirs, but also identified additional pathways that may become vulnerable in the virus-infected cellular state, such as metabolic and transport processes, autophagy, biogenesis and maintainance of organelles, and polymerase II mediated transcription, that may contain significant SL targets. Furthermore, our 738 high-confidence SL targets were enriched for multiple Reactome and GO terms associated with viral infection, confirming the relevance of the targets within the infection context. Our data suggest that inclusion of many diverse omics datasets as well as multiple CRISPR studies generates a large pool of SL candidate genes for further prioritization and validation.

Broadening our investigation, we identified SL targets that may be pan-viral. Even with their distinct life cycles and biology, both coronaviruses and influenza viruses shared common host SL targets. While only eleven of our 71 targets were enriched in virus-associated pathways, our data suggest that the number of pan-viral targets based on synthetic lethality might be substantially larger. One known example is 60S ribosomal protein L28 (RPL28) which plays a role in negatively regulating an influenza A virus encoded peptide for antigen presentation, thus potentially modulating immunosurveillance (85). Another candidate SL target, charged multivesicular body protein 2A (CHMP2A), is a member of the endosomal sorting complex required for transport (ESCRT)-III machinery (86) and interacts with SARS-CoV-2 Orf9b. Furthermore, CHMP2A has been shown to contribute to the budding of a variety of viruses, including HIV (87), equine infectious anemia virus (EIAV) 401, and murine leukemia virus (88), suggesting a critical role for virus release and thus a potential role as pan-viral SL target.

Thus, our discovery significantly widens the future exploration scope for therapeutic strategies against multiple viruses, using combined drug regimens that could target two or more host genes at otherwise subtherapeutic levels, potentially minimizing off-target effects.

It is important to note that our methodology, while promising, is not without limitations. We recognize that no single dataset or database can predict or validate virus-specific SL targets with absolute accuracy. For example, CRISPR KO depleted genes are not a true “gold standard” for SL prediction, but they currently provide the best available type of data suitable for assessing performance in support of a gene’s SL candidacy. Additionally, the SynLethDB database used here contains synthetic lethal gene pairs derived primarily from cancer cell lines and tumors (54). Thus, while it includes many SL interactions that are relevant in both cancer and viral infection states, it will contain some SL targets that are irrelevant in the virus-infected cellular state. It also may lack some targets that are important in viral infections. Yet, when available data are combined in the manner demonstrated in our study, they can predict a significant number of candidate SL genes that can be further constrained and validated in laboratory settings using virus-infected cell lines and animal models.

For example, to prioritize targets, fold-change values in omics (27)and CRISPR data sets could be used to rank virus-impacted host proteins (VIHs) and functional depletion, respectively. The “SL statistics” score provided by SynLethDB and the number of VIH partners paired with a predicted SL target (27) could provide additional measures. Arguments that have been shown to enrich for SL interactions in a more generic manner include pairing paralogs (84) as well as co-regulated or mutually exclusive gene pairs (89, 90), in addition to pairing genes that exhibit certain network topology features (91). Some of these data-driven algorithms are part of SL-Cloud (90), a new platform that was developed for the cancer field but is essentially agnostic and can be utilized for infectious disease research. Any of the above metrics alone or in combination could be integrated into our pipeline upon laboratory confirmation of their usefulness in the viral infection context, enhancing precision and virus-specificity.

To summarize, our research has demonstrated that virus-specific SL targets can be predicted from various omics data classes, with varying levels of predictive power. Furthermore, these SL genes are highly enriched in depleted CRISPR KO data from virus-infected cells. Thus, our pipeline allows for a new application of many already existing high-throughput infectious disease datasets which are often under-utilized. In human cells, each VIH was seen to form SL interactions with an average of three genes ; in yeast, every gene is in at least one synthetic lethal relationship across different growth conditions (10, 92). Hence, it was not surprising that our study found many candidate SL targets for the two viruses investigated, all having the potential to specifically kill virus-infected cells. These candidate SL targets add substantially to the number of potential host-based antiviral targets previously described in the literature. With further validation, our presented strategy for identifying SL targets has the potential to identify additional SL targets for other viruses and cell-dependent pathogens, leading to new specific and broadly-targeting host-based therapies against infectious diseases.

## DATA AVAILABILITY

No new data were generated or analyzed in support of this research. All available data used in this article are referenced in the text or tables and have been made public by the respective authors.

## SUPPLEMENTARY DATA

Supplementary Data are available at NAR online.

## AUTHOR CONTRIBUTIONS

J.P.S.: Conceptualization, formal analysis, methodology, visualization, writing—original draft. M.L.N.: Methodology, formal analysis, visualization, writing—review & editing. A.N.: Conceptualization, writing—review & editing. F.M.: Conceptualization, writing—review & editing. J.D.A.: Conceptualization, writing—review & editing.

## ACKNOWLEDGEMENTS

We thank Fergal J. Duffy for guidance on questions related to specific R packages. We thank Jason P. Wendler for guidance and helpful discussions related to the generation of the pipeline.

